# Capture-recapture for -omics data meta-analysis

**DOI:** 10.1101/2023.04.24.537481

**Authors:** Julius Juodakis

## Abstract

One of the major goals of modern -omics studies, in particular genome-wide association studies (GWASs), is to understand the polygenicity of various traits, i.e. the number of genetic factors causally determining them. Analogous measures could also be used to estimate the number of trait markers from non-genetic studies, such as proteomics or transcriptomics.

Here, we describe how capture-recapture (C-R) models, originating in animal ecology, can be applied to this task. Our approach works by comparing the lists of trait-associated genes (or other markers) from several studies. In contrast to existing methods, C-R is specifically designed to make use of heterogeneous input studies, differing in analysis methods, populations or other factors: it extrapolates from their variability to estimate how many causal genes still remain undetected.

We present a brief tutorial on C-R models, and demonstrate our proposed usage of it with code examples and simulations. We then apply it to GWASs and proteomic studies of preterm birth, a major clinical problem with largely unknown causes. The C-R estimates a relatively low number of causal genes for this trait, but many still undetected protein markers, suggesting that diverse environmentally-initiated pathways can lead to this clinical outcome.

## 1 Introduction

Many important human traits are determined by complex molecular networks that involve a large number of genes and proteins. There is considerable interest in measuring the extent of such a network for a chosen trait, or polygenicity in its broadest sense. This property is key for our understanding of the overall molecular control principles, for example, distinguishing between the hypotheses proposing modular [1], polygenic [2] or omnigenic with core genes [3] architectures. More pragmatically, polygenicity estimates can help in the search for disease causes or biomarkers: they can be used to tune study design for better power [4], or broadly indicate whether we should focus on discovering additional causal factors or more complex interactions of known ones [5]. While modern -omics experiments are designed to directly assess which genes, transcripts or proteins are involved in a particular biological process, differences between populations, study designs, and random variation mean that each experiment will still only identify some of the relevant factors, and statistical estimation of polygenicity is needed.

Currently, polygenicity is commonly estimated using results from genome-wide association studies (GWASs). The main approach is to model the effect sizes of genetic variants on the trait (obtained e.g. by regression) with a mixture distribution of lower-variance and higher-variance components, representing null and causal effects. The estimated weight of the causal component(s) is taken to measure the proportion of causal variants, and thus polygenicity at the variant level [4, 6–9]. Recently, this approach was also applied at the gene level, by first combining the variant effects into imputed transcript expression changes and fitting a mixture distribution to these [10]. Alternatively, polygenicity has been estimated using the kurtosis of the observed effect size distribution [11] or the kurtosis of gene-level heritabilities [12]. (The methods directly apply to the meta-analysis setting as well, using pooled effect sizes.) In short, the methods rely on a precisely chosen distribution of null effects, and infer polygenicity based on deviations from that.

Measures analogous to polygenicity could also be estimated from other types of -omics data. Expression microarrays are often used to infer various properties of the underlying genetic networks, such as connectivity [13]. We are not aware of any explicit estimators for network size from such data, but in meta-analyses, overlap between lists of differentially expressed genes from the studies is commonly reported, e.g. [14, 15], and various similarity measures for such lists have been developed [16, 17]. Meta-analyses in proteomics also typically report such overlaps of associated markers, e.g. [18–20]. Assuming the detected proteins or transcripts indeed represent causal genes, the amount of overlap between study results serves as a clue about the polygenicity of the trait.

While the mixture methods used in GWAS are well formalised statistically, they are not designed to use heterogeneous data. Specifically, we consider the problem of meta-analysing studies that are focused on the same trait, but differ in some biological or statistical design aspects, such as: experimental techniques; statistical models (e.g. binary or continuous analysis); aspects of the trait definition or inclusion criteria (e.g. earlier- or later-onset cases); sampling tissue and time point, for non-genetic studies; and others. Likely, different genes or biomarkers emerge in each setting. Our goal is not to average out their effects over the settings, as done in standard meta-analysis, but rather to use this variety to gain insights into the polygenic structure underlying the trait as a whole.

In this paper, we will show how the *capture-recapture* framework (C-R) can be applied to this problem, i.e. to estimate polygenicity from heterogeneous -omics data. C-R statistically formalises the idea of inspecting list overlaps: it originates from ecology, where it is used for estimating the total populations of animals by combining several capture attempts. We will present an introduction to C-R and how it applies to -omics, demonstrate the usage and performance of the relevant tools, and analyse two real-data problems of preterm birth.

## 2 The capture-recapture framework

### 2.1 The concept

To understand the intuition behind C-R, consider a field experiment consisting of two surveys, in which animals are captured within a defined area, such as a forest. In the first survey, *n*_1_ animals are randomly caught, then marked and released, mixing back into the population. After a period of time, a second survey is conducted, randomly catching *n*_2_ individuals, and *m*_2_ of these are seen to carry the mark from the first survey. Then, we can expect that the first survey marked a fraction of the population close to *m*_2_*/n*_2_, and so we have an intuitive way to estimate *N*, the total number of animals in the area: 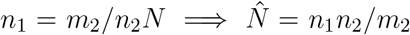; this is the Petersen-Lincoln estimator and the simplest case of C-R. (For in-depth technical details and historical references of the models discussed here we refer the reader to [21–23].)

Throughout the last 100 years, the C-R idea has been formalised and extended to accommodate a wide variety of data. The concept is not in any way specific to ecological surveys or physical capturing, and can be applied to any situation where a population size needs to be estimated from multiple lists of detected individuals. C-R is commonly used in demographics and public health, e.g. for estimating disease prevalence by combining sources such as hospital records, prescription registers, and insurance claims [21, 24], and especially in situations where direct counting is difficult, such as stigmatising or criminal outcomes [25, 26].

### 2.2 Model details

More precisely, the input data in C-R settings is the capture history for each observed individual, such as “1101…”, with 1 at position *i* indicating that the individual was observed on the *i*th occasion and 0 that he was not. We will denote by *n*_1101…_ the count of such a capture history; for example, for a three-list dataset, we have *n*_100_ – the number of individuals recorded only in list 1, *n*_010_ – the number of individuals recorded only in list 2, *n*_110_ – the number of individuals present in both lists 1 and 2, but not 3, etc. We seek to estimate *N* = *n*_000_ + *n*_100_ + *n*_010_ + *n*_001_ + *n*_110_ + *n*_101_ + *n*_011_ + *n*_111_, or just the *n*_000_, the number of individuals that were not seen in the dataset.

Since practical applications of C-R often involve more than 2 lists, solving the model requires more complex techniques than the Petersen-Lincoln estimator. (We will continue assuming that 3 lists are used, to keep notation clearer.) Several approaches are common. One is to define a statistical model of the capture process, with constraints based on the domain knowledge, and estimate *N* by maximising the probability of observed data (likelihood). Darroch, Otis and Pollock developed a series of models in this fashion [27]. The simplest, **M0**, assumes that each individual in each list is detected with a fixed probability *p*, and independently of others. Then the likelihood can be obtained from the multinomial distribution as:

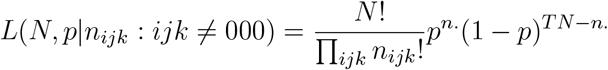

where *n*. is the total number of recorded detections in the dataset, *T* is the number of lists. Note that maximising the likelihood also provides an estimate of *p*, which may be of interest too. If the detection probability changes between lists, we have model **Mt**, with likelihood [27]:

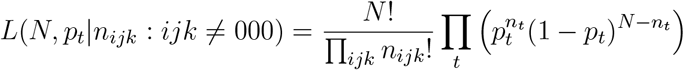

with *n*_*t*_ – the number of detections in the list *t*. Similarly, heterogeneity between the individuals can be incorporated, as a unique *p* for each individual (model **Mh**). There are also models accounting for behavioural response to capture (**Mb**) and combined extensions (**Mtb** etc.), which are more relevant in ecological applications. Overall, extensibility is one advantage of the likelihood approach, and it can easily incorporate any additional information recorded with the detection, for example, if some measure of the detected signal strength is available [28].

An alternative solution for multiple lists is to create a log-linear model for the counts of each type of capture history. For three lists, one such possible setup is: log *E*(*n*_*ijk*_) = *u* + *iu*_1_ + *ju*_2_ + *ku*_3_ + *iju*_12_ + *jku*_23_ + *iku*_13_ [24, 29]; estimating *u* = log *E*(*n*_000_) is of interest and allows us to estimate *N*. The terms allow different main effects for each survey (in this example, *u*_1_ are the log-odds for being detected in list 1 etc.) and any pairwise or even higher interactions between the lists, so models **M0, Mt** and certain others can be obtained in this approach as special cases. Common GLM software can be used for fitting. This is also the approach taken in the R package Rcapture, which we will use in this paper [30].

Note that all the models discussed here share the assumption that the population to be estimated is “closed” – no individuals enter or leave it between the surveys. This can be a major limitation in certain ecological contexts, and much work has been done on adapting C-R to include births or deaths. As we will deal exclusively with closed populations in this paper, we do not discuss these extensions further and refer the interested reader to [23].

### 2.3 Adapting C-R for -omics

Our premise is to directly apply the C-R models to the -omics meta-analysis setting. Thus, the “captured” objects are genes (or other biomarkers) causing a trait of interest; the population to be estimated, *N*, is the total number of causal genes for that trait; the observed data are lists of associated genes, retrieved from 2 or more studies of the trait. In effect, applying C-R will allow us to formally measure the overlap between the lists, and provide an estimate of the number of causal genes missed in both studies, as well as the total number of causal genes for this trait (polygenicity).

We assume that the included studies can detect different genes. Several reasons commonly lead to this in practice [31–33]: random variation between the samples; true differences in effect sizes between studied populations; differences in study design and technical methods; different inclusion criteria, covariate adjustment, trait definition, etc. We will highlight some specific instances of such factors in our case studies further below. In the C-R approach, the detection process is treated as a black box – the model simply includes a detection probability term. Based on the variability between the studies’ results, C-R estimates both this probability and the number of genes remaining undetected.

Specifically, we will consider these models:

- **M0**. Each gene is detected in each study with equal probability *p* (no heterogeneity, no study effects).
- **Mt**. In study *t*, each gene is detected with a probability *p*_*t*_ (study effects).
- **Mh**. Gene *g* is detected in each study with a probability *p*_*g*_ (heterogeneity between genes). We will specifically consider the model **Mh Poisson2** where the *p*_*g*_’s are based on a mixed Poisson distribution [30].
- **Mth Poisson2**. In study *t*, gene *g* is detected with a probability *p*_*t,g*_ (heterogeneity between genes and study effects), based again on the Poisson mixture.

The latter models are likely more realistic – we expect some studies to have higher detection power, e.g. due to sample size, and certain genes likely have higher effects in all contexts and thus are detected often. However, fitting models with heterogeneity requires more data, while ignoring these differences generally leads to underestimated *N*, so the simplified models still provide valid lower bounds [23].

Key assumptions are that the total “population” of causal genes is closed (does not change between the studies), as discussed before, and that the genes are reliably identified by their names. These assumptions are violated e.g. if some of the input studies only analyze certain chromosomes or subsets of proteins. The analyst also has to ensure that the protein names or other identifiers used are consistent across the studies. We also do not expect the genes to “react” to a detection in any way, i.e. the detection probability does not depend on the temporal order of the studies. With these assumptions, we avoid the need for the more advanced C-R models developed for other areas [21, 23].

### 2.4 Implementation and example code

We fit the C-R models using the R package `Rcapture` [30]. The package takes as input a detection history, sets up a log-linear (Poisson) regression based on the tested model, and solves the model to produce the abundance estimate. Several different models are included in `Rcapture` (although given 2 lists, only **M0, Mt** and **Mb** can be fitted).

As the starting input, we simply create a list recording the detections of each study in a separate element. For example:

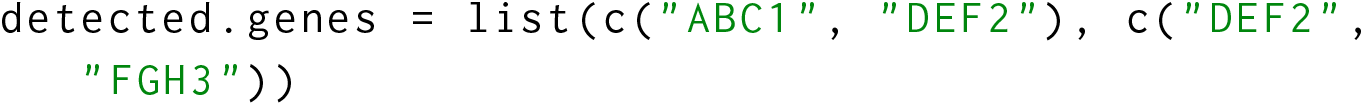

The list is converted into a capture history compatible with Rcapture, and the available models are fitted, with these three lines of code:

~~~
all. genes = unique (unlist (detected. genes))
capt. hist = sapply (detected. genes, function (x)
 all. genes % in% x)
closedp (capt. hist)
~~~

This example analysis would produce the output below, showing 4 as the estimated number of causal genes (“abundance”); this matches the simplest 2-list estimator 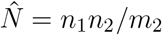. Akaike information criterion (AIC) values are also reported and can be used for model selection. Note the `infoFit` column which here warns about a bad model fit, due to the very low amount of data in this example.

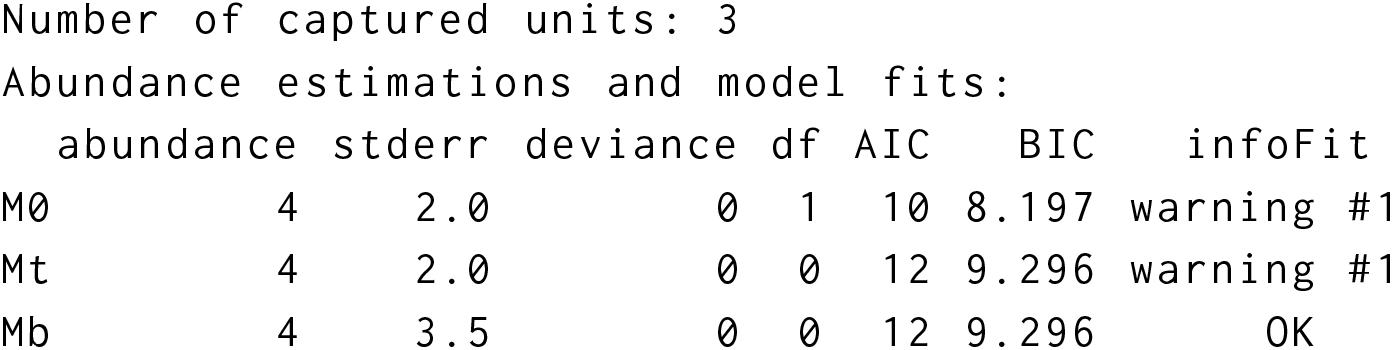

Confidence intervals can be calculated from the provided standard errors, i.e. 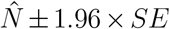 for 95 %, or better ones (based on profile likelihood [30]) can be obtained using the `closedpCI.*` functions with the desired model:

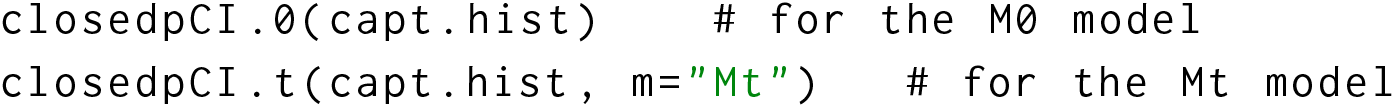

## 3 Simulations

### 3.1 Methods

We conducted simulations to verify that the proposed framework can recover the number of causal genes (or other biomarkers). First, we assumed a study design where the genes are directly identified. From a list of 2552 genes comprising human chromosome 1, we randomly assigned 30 genes as causal, and simulated 2 studies, each having a 10 % probability to detect each causal gene (i.e. each study has 10 % power, or sensitivity, for each gene). The two resulting lists of “detected” genes were analysed with C-R, model **M0**, implemented in `Rcapture` [30]. The process was repeated with different numbers of causal genes and detection probabilities, in 300 replicates in each setting.

Next, we simulated a meta-analysis of 2 GWAS studies, including SNP-level effects. For each study, we simulated the genotypes of 2000 individuals at *≈* 90, 000 positions (*X*) with the R `sim1000G` package [34]. As the reference (`sim1000G` uses it to determine the linkage structure and allele frequencies), we used genotypes of chromosome 1 from the 1000 Genomes European (CEU) samples, selected to have minor allele frequency in Europeans *>*0.05 and then downsampled to approx. 90,000 markers. The genotypes were created once and used in all simulation replicates. We then randomly selected 50 genes to be causal. For each study and each gene, a random SNP *s* within 10 kbp of the gene was selected, and assigned an effect size of *β*_*s*_ *∼* Uniform(*−*10, 10). A continuous phenotype was defined:

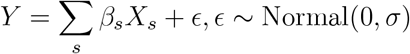

with different values of the residual standard deviation *σ*. The resulting heritability (*V ar β*_*s*_*X*_*s*_*/V ar Y*) is in the range of 25–95 %, relatively high to allow simulating fewer individuals than typical in GWAS.

A GWAS was conducted on this with PLINK 1.9 [35], using linear regression of the minor allele dosage as the association model. SNPs significant at the genome-wide threshold *p <* 5 *×* 10^*−*8^ were assigned to the closest gene (counting from the gene midpoint). The resulting gene lists from the 2 studies were analysed with the C-R model **M0**, as described above. We also repeated the C-R analysis using only genes tagged by independent SNPs (*r*^2^ *<* 0.1 if within 250 kbp, and *p <* 5 *×*10^*−*8^), selected with PLINK `--clump` command. Another set of simulations was carried out identically, but with 200 causal genes, and the SNP effect drawn as *β*_*s*_ Normal(0, 5). The simulations were repeated 300 times in each setting.

A third set of simulations was carried out as above, but using 3 GWAS studies with 1000, 3000 and 9000 individuals. Genotypes and phenotypes were simulated as above, again in two scenarios: 50 causal genes and a SNP effect drawn from Uniform(*−* 10, 10), or 200 causal genes and effect *∼* Normal(0, 5), residual SD *σ* set to 50. Association tests were conducted, significant SNPs clumped and assigned to genes as above. Gene lists were analysed with C-R models **M0, Mt** or **Mth Poisson2** in the Rcapture package.

### 3.2 Results

We verified that the C-R model is a consistent estimator for the causal gene number in simple situations where genes are identified directly in the studies (Figure 1). Since it requires at least 1 overlapping record between the studies, it fails when both the number of genes and detection probability are low, but in our simulations even *≈* 10 detected genes per study was enough to achieve useful estimates (Figure 1).

**Fig. 1.**
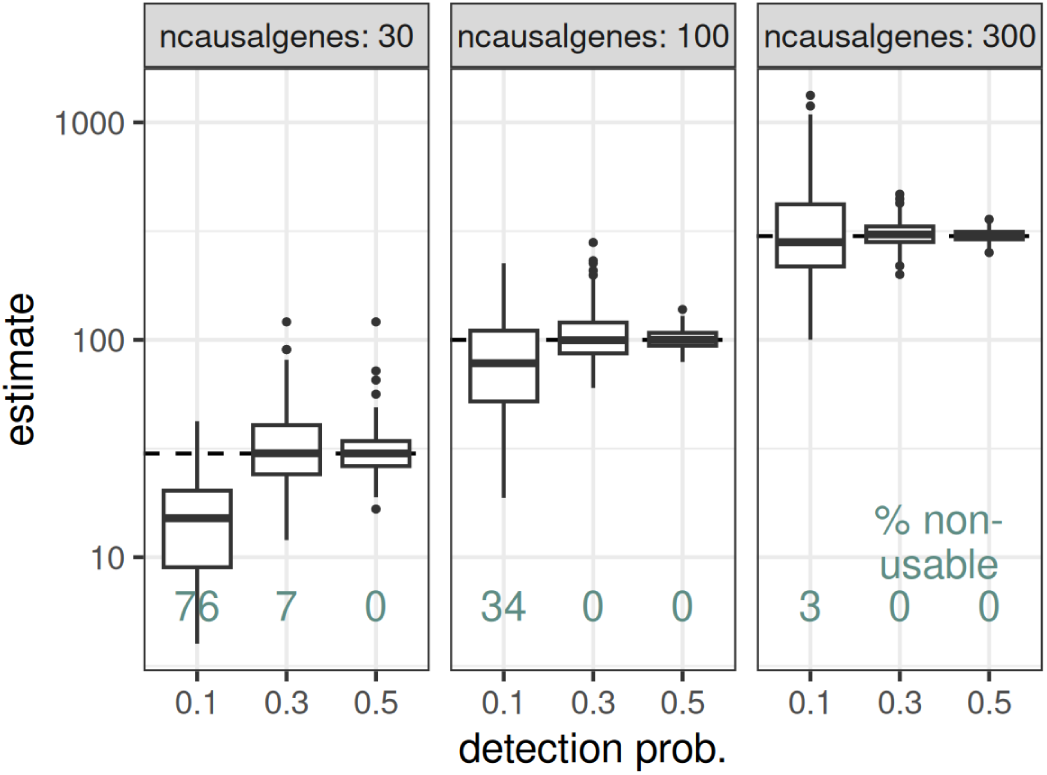
Distribution of the C-R (model M0) estimated numbers of causal genes, in simulations of pairs of studies directly detecting genes. In each simulated study, genes were directly drawn from the causal set, with the detection probability for each gene as shown on the x-axis. Also shown is the percentage of simulations that were not usable in C-R, i.e. the studies had zero overlaps.

When simulating GWAS studies, the problem of assigning SNPs to the correct causal gene is introduced. The estimate from C-R can be biased upwards, if the LD structure is ignored, because the causal signal spreads between linked SNPs which can then tag several different genes (Figure 2). However, in practice, researchers would assign such SNPs to the same gene, after standard post-GWAS procedures such as inspecting the linkage structure or conditional analysis; to approximately emulate this, we clumped the SNPs to an independent set of signals, and the bias mostly disappeared (Figure 2). We expect the bias to decrease further as researchers tend to define causal loci consistently with previous studies, which we did not model. Note also that even in the high-noise settings when up to half of the simulated meta-analyses fail, the remaining estimates are still roughly accurate (Figure 2).

**Fig. 2.**
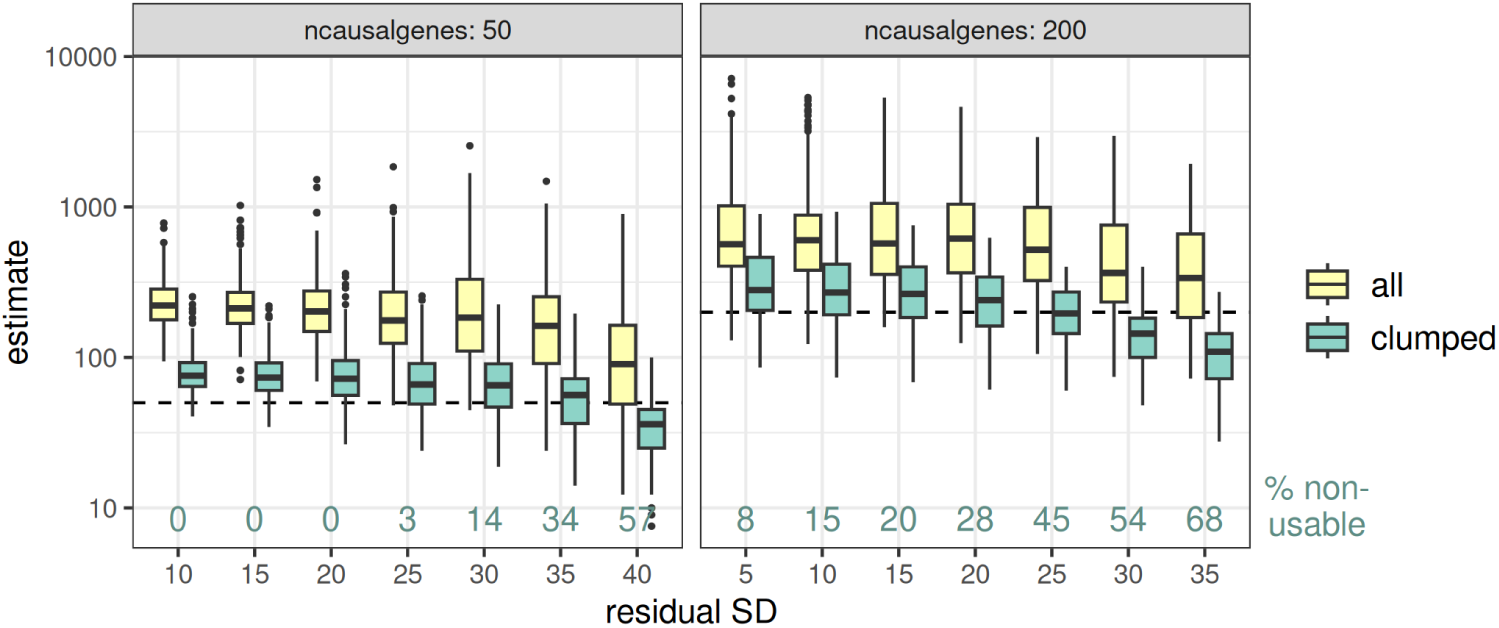
Distributions of the C-R (model M0) estimated numbers of causal genes, in simulations of pairs of GWAS studies. Each study simulates 90,000 SNPs for 2000 individuals, a phenotype with the specified residual standard deviation (higher SD = lower heritability), tests the SNPs for association with the phenotype, and reports genes corresponding to all associated SNPs (“all”) or only independent signals (“clumped”). Reported genes are analysed with C-R. Also shown is the percentage of simulations that were not usable in C-R, i.e. the studies had zero overlaps.

With 3 or more studies, other C-R models in addition to **M0** can be applied. We tested this on simulations of triplets of studies, differing in sample size, and thus the power or probability to detect the causal genes. Under this heterogeneity, model **M0** was substantially biased upwards (Figure 3). This was mostly fixed with the appropriate model **Mt** (Figure 3), again with a small bias remaining due to assigning linked SNPs to multiple genes, which would be resolved in practice. (About 7–10 % of the simulations had no overlap between the studies and could not be analysed.) We also observe that including a between-gene heterogeneity component (model **Mth Poisson2**), which was not needed here, can greatly reduce estimator precision.

**Fig. 3.**
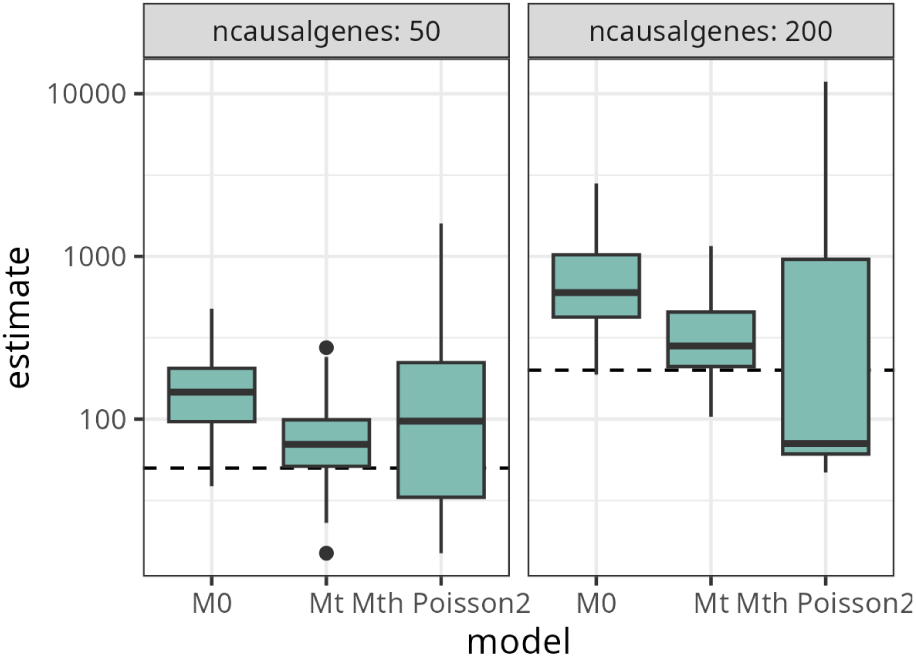
Distributions of the C-R estimated numbers of causal genes, in simulations of triplets of GWAS studies. In the studies, 90,000 SNPs are simulated for 1000, 3000 and 9000 individuals, then tested for association with a simulated phenotype, and genes corresponding to the independently associated SNPs are reported. Reported genes are analysed with different C-R models.

## 4 Case studies

To demonstrate how the proposed method can be applied to real problems, we will use it to analyse different -omic studies of human pregnancy duration. Deviations from the expected duration of 40 weeks, in particular preterm birth (birth *<*37 weeks), have major consequences to neonatal survival and health [36]. Several genome-wide association studies of pregnancy duration have been conducted [37–39], as well as many proteomic, transcriptomic and candidate-gene studies attempting to identify any relevant biomarkers. Still, a large fraction of the variability in this trait remains unexplained. It is considered that pregnancy timing control is very heterogeneous, with multiple causal pathways leading to apparently similar outcomes [40]; accordingly, studies have used various different case definitions or statistical models, with varying success. Using C-R, we aim to summarise these results, and give an approximate indication of how many causal genes and proteins still remain unseen in the current studies.

### 4.1 GWAS of preterm birth and gestational age

Two large maternal GWASs of pregnancy duration have been conducted recently [38, 39]. Both studies analysed the samples in two statistical models: case-control tests with preterm delivery as the outcome, and regression models of continuous gestational duration. Still other models have been proposed and used in the field, such as testing post-term vs. term pregnancies, more extreme pretermity cutoffs, survival and rank-transformed models. To estimate how many genes remain undetected by the two models used so far, we applied C-R to the results of the two studies.

From each study, we retrieved two lists of genes, i.e. associations with PTD and with GA, as reported by the authors (generally, for each independently genome-wide associated SNP the nearest gene was chosen). In Solé-Navais et al. [38], 23 genes were reported for GA, and 7 for PTD, of which all but 1 were also reported for GA. The Pasanen et al. [39] study reported 15 genes for GA and 4 for PTD (of these 2 unique to PTD). Some samples were used in both studies, and the top hits were largely shared, but 14 of the genes were still unique to [38] and 7 unique to [39].

We applied C-R models to the pair of lists in each study (Table 1). The estimated number of genes was around 30 in each (model **Mt**). This is not much higher than the numbers of genes detected directly – corresponding to the high overlap observed between the GA and PTD lists – and indicates that analysing this data with furhter association models (such as the post-term or survival ones) likely would not reveal many more genes. (The **M0** models are less appropriate here as they assume equal detection power in both lists, and also showed higher AIC.)

**Table 1.** C-R estimates of the number of causal genes, based on different C-R models and input studies. From each study, 2 gene lists are retrieved, corresponding to gestational duration and preterm delivery analyses.

| Gene lists | model | estimate | SE | AIC |
| --- | --- | --- | --- | --- |
| Solé-Navais et al. [38] | M0 | 37.5 | 9.2 | 31.6 |
| Solé-Navais et al. [38] | Mt | 26.8 | 3.6 | 16.3 |
| Pasanen et al. [39] | M0 | 45.1 | 25.2 | 22.7 |
| Pasanen et al. [39] | Mt | 30.0 | 14.0 | 15.6 |
| both studies | M0 | 40.7 | 5.3 | 73.0 |
| both studies | Mt | 37.0 | 3.9 | 52.1 |
| both studies | Mh Poisson2 | 65.6 | 23.1 | 68.2 |
| both studies | Mth Poisson2 | 58.1 | 18.9 | 43.3 |

Note that the C-R estimates can only account for the sources of variability reflected in the input data. In this example, the two gene lists were produced by different analysis methods, but in the same cohorts. Thus, the estimates do not capture, for example, possible variation of effect sizes between populations. To incorporate that, the C-R models can also be applied to all four gene lists from the two studies. The estimates increase to around 37–66, depending on the chosen model (Table 1; as more than 2 lists are present, **Mth** models can also be fitted here). In other words, we can expect that, in addition to the ≈ 30 hits reported from these GWASs so far, at least around 10–30 more could still be discovered by conducting additional GWASs in other populations. (Noting that exceptionally large studies may find many more hits by capturing genes with very small effects, which are not reflected in any of the current GWASs.)

### 4.2 Proteomics of preterm birth

In the next example, we apply C-R to a larger set of studies which differ in various clinical and technical aspects. We conducted a literature search and meta-analysis of proteomic studies of preterm birth biomarkers. Studies were identified from the Web of Science database on 2022 August, using the following search string: `proteom* AND preterm AND (birth OR delivery)` in abstract. The search produced 164 abstracts, which were then manually inspected to remove studies of cell cultures or other organisms, studies of outcomes not primarily related to birth timing, reviews or other studies not reporting new results, as well as targeted multiplex or microarray studies (such designs would prevent some markers from being detected and so are not compatible with our method); 47 abstracts passed these criteria.

False positive detections are not accounted for in our method and would inflate the estimates upwards. To reduce their rate, we required the protein detections to pass a strict false discovery rate (FDR) criterion of 5 %. This meant that studies were excluded if they did not report any significance metrics, did not provide sufficient information to understand their metrics or calculate FDR, or if no significant proteins remained after authors’ adjustments or validation. Note that the method in general does not require significance metrics in the input, as the user only needs to provide the (reliably) detected marker names. After the final exclusions, 12 datasets from 10 studies remained, with details shown in Supplementary Table S1 [20, 41–49]. The studies cover a range of sampling time points, inclusion criteria, different protein identification techniques, different tissues and fractions (maternal plasma, amniotic fluid, placental tissues); although we excluded several studies which collected neonatal samples after birth, reasoning that these likely reflect consequences rather than causes of preterm delivery. In sum, these studies reported 385 detections (at our threshold of FDR *<*5 %), covering 311 unique proteins. Figure 4 shows the overlap between the 4 studies with the most detections: clearly, some of the proteins were common between multiple studies, but it is difficult to make any quantitative conclusion from such visual inspection, especially given that several more studies are not shown.

**Fig. 4.**
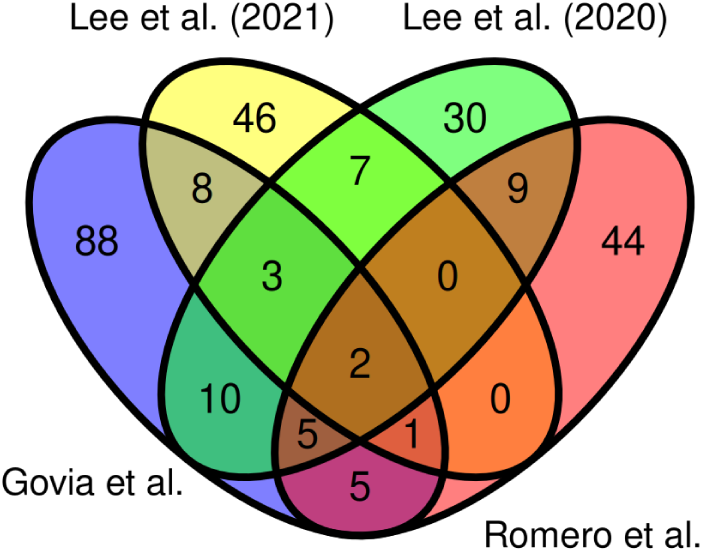
Venn diagram showing the overlap between the proteins reported in the 4 studies with the most detections from the meta-analysis of preterm birth biomarkers.

In contrast, the C-R models quantify the overlap, and directly produce an estimate of the total number of preterm birth biomarkers: 832 (95 % profile-likelihood CI 689–1030) using the **M0** model, or 757 (95 % CI 631–930) using the **Mt** model. The latter is likely more accurate as it includes a component to model the substantial heterogeneity present between studies, and is also favoured by the data, as reflected in the much lower AIC (203 for **Mt** vs. 550 for **M0**). Including between-gene heterogeneity components (**Mth** models) provided virtually no improvement (AIC 194–201). As the estimated number of markers is large and several times higher than detected in any single study in this review, we conclude that the proteomic signatures of preterm birth are highly diverse, representing different, potentially environment- or ancestry-dependent, etiologies.

## 5 Discussion

Capture-recapture modelling has found useful applications in many fields. In this paper, we showed how it can be applied to -omics as well, specifically when heterogeneous data sources are meta-analysed. In the case study of preterm birth, C-R indicated a relatively small number of causal genes and high overlap between GWASs with different analytic approaches, and in contrast estimated many biomarkers at the protein level. Our interpretation is that diverse environmental pathways lead to preterm birth, in line with e.g. [40], while the main genetic pathways to this outcome are few, but robust across samples and conditions.

While C-R models can be straightforward to apply, interpreting the results requires great care. Firstly, any biases present in the input data remain, in particular the potential for confounding. At the transcript or protein level, many of the detected changes might reflect consequences of the trait, not its causes [50]. Genetic variant associations are robust to many types of confounding, but even then it may still arise e.g. due to participation bias [51]. In our proteomics case study, we only include studies where the samples are taken before birth, to reduce such issues, but still the detection lists – and our estimates – may include many proteins which are non-causal biomarkers.

Secondly, since C-R works by extrapolating from the variability in the input studies, it can only account for the sources of variability that were present there. In particular, if all input studies were conducted on European individuals, the estimated polygenicity may miss some causal factors that are important in other populations, and will likely be too low. Thus, the usefulness of our method critically depends on the availability of relevant and diverse input data.

Finally, we emphasise again that polygenicity in our framework is defined differently than in the current alternatives. In C-R, we expect that the causal factors must become apparent and detectable in at least one testable setting.

In contrast, mixture methods such as [8] define polygenicity as the weight of the higher-variance component of the effect size distribution – which means that most of the causal factors are in fact modelled to have zero or near-zero effect. One could, in principle, meta-analyse different studies and apply the latter methods to the distribution of averaged effect sizes. However, even simple differences in design – such as binary vs. continuous outcome – lead to complications when averaging the effects [52]. Harmonising effect sizes for meta-analysis can also be a challenge with expression arrays [53]. We therefore see C-R, mixture models, as well as other methods for quantifying heritability and genetic correlation [54] as complementary approaches, and combining and contrasting these will be needed to fully understand the challenges of human complex trait genetics.

## Acknowledgements

The study was made possible by funding from Lilla Barnets foundation awarded to the author, and support from Swedish Research Council (2019-01004) and Agreement concerning research and education of doctors (ALFGBG-965353) grants awarded to Bo Jacobsson. We are grateful to all the participants of studies that were reviewed and re-analysed here.

## Data availability

The analysis code and input datasets used here are available at https://github.com/jjuod/geneCR.

