## Supplementary Material for "Capture-recapture for -omics data meta-analysis"

### 1 Supplementary Material

| ID and reference | Setting | Method | Proteins quantified | Hits at 5 % FDR |
|---|---|---|---|---|
| 1. Huang et al (2020) | sPTD cases (n=15) and matched term controls (n=15), serum at 15–20 wk | LC-MS | 460 | 0 |
| 3a. Tiensuu et al (2022) | sPTD <35 wk cases (6), full term controls (6), placenta basal plate biopsy | 2D-DIGE-MS | > 200 | 0 |
| 3b. | As above, but chorionic plates | As above | > 200 | 0 |
| 6. Kim et al (2021) | sPTD cases (5), term controls (20), cervicovaginal fluid at <20 wk | LC-MS | 990 | 52 |
| 7. Lee et al (2021) | Plasma from threatening PTD patients who delivered sPTD (10) vs who delivered at term (10), matched, without IAI | LC-MS | 244 | 71 |
| 10.Hong et al (2020) | Patients with PTL <33 wk, without intra-uterine infection/inflammation, sPTD <34 wk cases (10) vs $\geq$ 34 wk controls (10), AF from amniocentesis | LC-MS | 780 | 2 |
| 11a. Dixon et al (2018) | Exosomes from AF samples collected during labour, sPTB <37 wk labour cases (13) vs term labour controls (11) | PAGE-MS | 501 | 2 |
| 11b. | As above, pPROM cases (8) vs term labour controls (11) | As above | 501 | 1 |
| 12. Lee et al (2020) | Patients with emergency cervical cerclage, sPTD<34 wk cases (8) vs $\geq$ 34 wk controls (8), AF from amniocentesis before cerclage | LC-MS | 624 | 68 |
| 13. Dan et al (2021) | Patients with emergency cervical cerclage <25 wk, maternal plasma samples, sPTD <33 wk cases (10) vs $\geq$ 33 wk controls (10) | LC-MS | 777 | 0 |
| 14. Romero et al (2009) | AF from patients with PTL, sPTD with IAI cases (24), vs controls delivered at term without IAI (26) | LC-MS | 325 | 67 |
| 15. Govia et al (2018) | AF from amniocentesis, cases with threatening PTL $\approx$ 22 wk (10) vs controls with routine amniocentesis (22) | LC-MS | 507 | 123 |

**Table S1** Details of the proteomic studies analysed in the presented case study.
